# Regulation of cellular LDL uptake by *PROX1* and *CHD7*

**DOI:** 10.1101/2022.09.20.507601

**Authors:** Candilianne Serrano-Zayas, Matthew L. Holding, Taslima G. Khan, Vi T. Tang, Jennifer M Skidmore, Donna M Martin, David Ginsburg, Brian T. Emmer

## Abstract

An elevated level of low-density lipoprotein (LDL) in the bloodstream is a causal risk factor for atherosclerotic cardiovascular disease (ASCVD). The low-density lipoprotein receptor (LDLR) is a critical regulator of circulating LDL, and increasing LDLR activity is an effective therapeutic approach to reduce circulating LDL cholesterol levels. In this study, we characterize *PROX1* and *CHD7*, two genes we previously identified in a genome-scale CRISPR screen as positive regulators of LDL uptake in HuH7 cells. We found that although disruption of either *PROX1* or *CHD7* significantly reduced LDL uptake, only *PROX1* disruption significantly reduced the cellular levels of *LDLR* mRNA and surface-displayed LDLR protein. Consistent with a direct role for *PROX1* in *LDLR* gene regulation, we also observed in publicly available data sets the presence of two liver-specific PROX1 binding sites near the *LDLR* locus, one of which colocalized with biochemical hallmarks of enhancer activity in hepatic tissue. Both PROX1 *LDLR* binding sites contained predicted PROX1 transcription factor binding motifs and colocalized with binding sites for HNF4α, a known interactor for PROX1 and regulator of hepatic lipid metabolism and LDL uptake. In contrast to PROX1, no CHD7 binding sites were detected near the *LDLR* locus. Together, our results support a model in which both *PROX1* and *CHD7* promote cellular LDL uptake through distinct mechanisms, with *PROX1* directly promoting *LDLR* gene expression and *CHD7* functioning through an LDLR-independent pathway.

## Introduction

Atherosclerosis is characterized by the accumulation of cholesterol-rich plaques in the arterial vessel wall, which may lead to progressive obstruction of the blood vessel lumen or acutely rupture and serve as a nidus for thrombus formation. Despite advances in treatment, atherosclerotic cardiovascular diseases remain the leading cause of global morbidity and mortality [1]. Evidence from diverse fields has firmly established elevated plasma low-density lipoprotein (LDL) as a causal risk factor in the development of atherosclerosis [2]. The concentration of LDL particles in the bloodstream is regulated by the hepatic LDL receptor (LDLR), which specifically binds the apolipoprotein B component of LDL particles. This binding triggers clathrin-mediated endocytosis of the complex into hepatocytes, removing the LDL particle from circulation [3]. Loss-of-function mutations in *LDLR* cause familial hypercholesterolemia, an autosomal dominant disease characterized by marked elevation in plasma LDL cholesterol and early onset ASCVD, while common genetic variants in putative LDLR regulatory regions are also associated with ASCVD risk [4]. Similarly, therapeutic agents that increase LDLR activity (e.g. statins, PCSK9 inhibitors) are effective for the prevention and treatment of ASCVD [5, 6]. Taken together, these observations support the central importance of hepatic LDL clearance in ASCVD pathogenesis.

In order to identify novel regulators of LDL uptake by hepatocytes, we previously performed a genome-wide CRISPR screen in HuH7 cells [7]. In addition to identifying a number of well-characterized LDLR regulators such as *LDLR* itself, *MYLIP, SCAP, MBTPS1*, and *MBTPS2* [3, 8, 9], this screen also detected several genes that did not have a previously established role in LDL uptake, including *PROX1*, encoding Prospero homeobox protein 1, and *CHD7*, encoding chromodomain helicase DNA binding protein 7. PROX1 is a transcription factor that has been implicated in a variety of developmental pathways [10–12]. CHD7 is a chromatin-modifying enzyme, and mutations in this gene result in an autosomal dominant developmental disorder known as CHARGE (<u>C</u>oloboma, <u>H</u>eart defect, <u>A</u>tresia choanae, growth <u>R</u>etardation, <u>G</u>enital abnormality, and <u>E</u>ar abnormality) syndrome [13–15]. We now report our characterization of *PROX1* and *CHD7* function in the regulation of cellular LDL uptake. Our findings confirm a role for both genes in promoting LDL uptake, and suggest that *PROX1* directly regulates *LDLR* gene expression whereas *CHD7* operates through an LDLR-independent mechanism.

## Results

### Cell density-dependent regulation of LDLR

Our initial attempts to validate the functional influence of *CHD7* and *PROX1* on LDL uptake were confounded by a high degree of variability between biological replicates. Given the previously reported influence of cell density on LDL uptake [16–18], we therefore adapted our flow cytometry approach to account for cell number, spiking in flow cytometry beads of a known concentration to extrapolate HuH7 cell densities and examine their relationship to cellular LDL uptake and LDLR expression. We found a clear negative correlation between cell density and both LDL uptake (Fig 1A) and surface LDLR abundance (Fig 1B). Interestingly, we observed an opposite correlation for *LDLR* mRNA (Fig 1C) and total cellular protein levels (Fig 1D). In subsequent experiments, we therefore quantified LDL uptake and LDLR expression for control cells across a range of plating densities and analyzed the effect of CRISPR targeting relative to this standard curve. This approach reduced experimental variability across independent biological replicates (Fig 1E).

**Figure 1.**
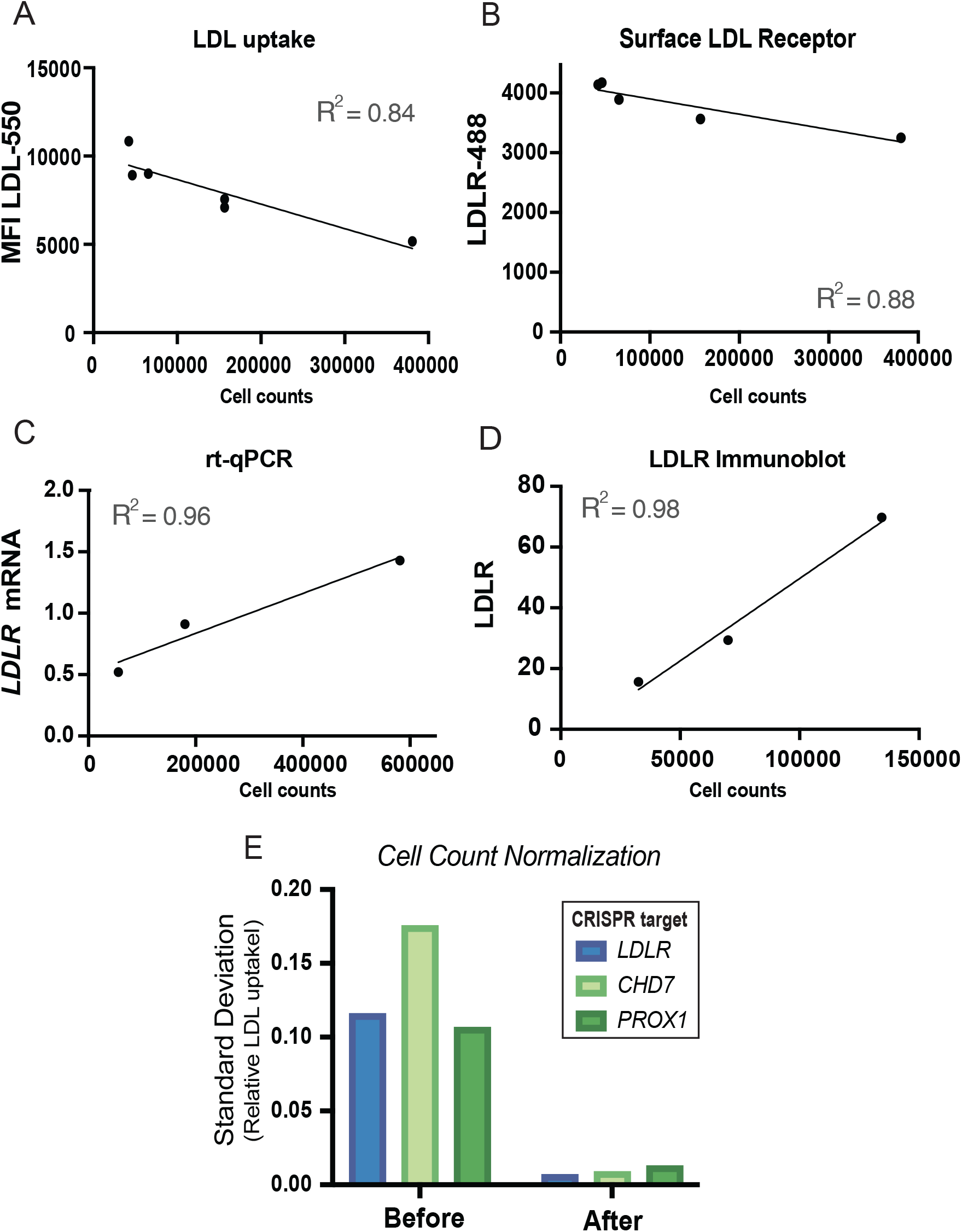
Density-dependent regulation of LDLR expression. Linear regression analysis of nontargeting gRNA-treated HuH7 cells plated in 6-well plates at a range of densities, with total cell numbers calculated based on the relative frequencies of cell events and bead events, with the latter occurring at a known density. The relationship of cell number at the time of the assay is plotted relative to quantification of (A) cellular LDL uptake by flow cytometry, (B) surface LDLR abundance by flow cytometry, (C) cellular *LDLR* mRNA levels as determined by rt-qPCR, and (D) total cellular LDLR protein abundance detected by immunoblotting. (E) Standard deviation of relative LDL uptake for each indicated CRISPR mutant for independent biological replicates before and after normalization to a standard curve of control cells plated at a range of densities.

### *PROX1* and *CHD7* promote cellular LDL uptake

We first examined individual gRNA enrichment for each of the 15 gRNAs targeting *PROX1* and *CHD7* in our previously published CRISPR screens of LDL uptake and LDLR abundance [7]. For both genes, depletion of nearly all individual gRNAs was observed in LDL^high^ cells relative to LDL^low^ cells, consistent with disruption of these genes reducing LDL uptake (Fig S1A). Interestingly, preincubation of cells in lipoprotein-depleted media, which stimulates upregulation of LDLR expression, led to a more pronounced depletion of *PROX1*-targeting gRNAs in LDL^high^ cells, while depletion of *CHD7*-targeting gRNAs was similar with or without preincubation in lipoprotein-depleted media (Fig S1B). *PROX1*-targeting gRNAs were also more clearly depleted than *CHD7*-targeting gRNAs in cells with high surface LDLR expression (Fig S1C). *CHD7-*targeting gRNAs but not *PROX1*-targeting gRNAs were depleted in LDL^high^ cells when screened on a *LDLR* knockout genetic background (Fig S1D). Together, these patterns of enrichment suggested that although both *PROX1* and *CHD7* appeared to influence LDL uptake, they may differ in their functional interaction with *LDLR*.

To validate the functional influence of *PROX1* and *CHD7* on LDL uptake and to examine their mechanism of action, we generated lentiviral CRISPR constructs harboring a single gRNA targeting each gene, selected based on its clear enrichment across independent screens in our previous report [7]. We transduced HuH7 cells with these constructs and verified depletion of each corresponding protein (Fig 2A-B). Flow cytometry analysis of CRISPR-targeted cells (with adjustment for cell density as noted above) confirmed a significant but partial decrease in LDL uptake resulting from CRISPR targeting of either *PROX1* (38% reduction, 95% CI = 20%-55%) or *CHD7* (25% reduction, 95% CI = 8%-43%) (Fig 2C).

**Figure 2.**
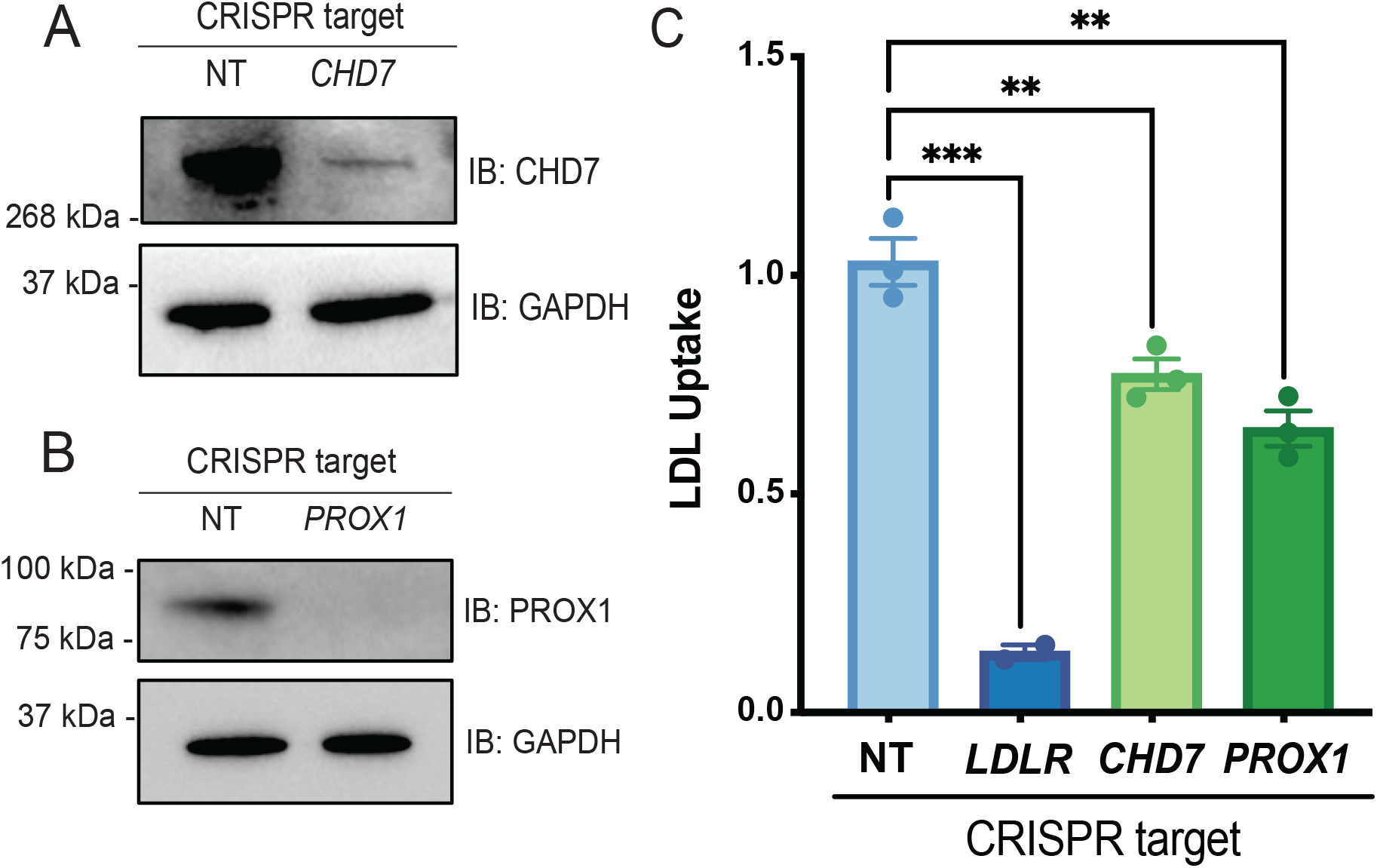
CRISPR targeting of either *CHD7* or *PROX1* decreases LDL uptake by HuH7 cells. (A, B) Immunoblot of CHD7 (A) and PROX1 (B) protein levels in non-targeting (NT), and *CHD7* CRISPR-targeted (A) or *PROX1* targeted (B) HuH7 cells, with lower panels showing an immunoblot for GAPDH as control. (C) Flow cytometry analysis of LDL uptake in *LDLR, CHD7* or *PROX1* CRISPR-targeted cells normalized to cells treated with a NT gRNA. Data points represent independent biological replicates (n=3), and error bars represent SEM (One-way ANOVA) (** p < 0.01, *** p <0.0001; each data point represents an independent biological replicate).

### Differential influence of *PROX1* and *CHD7* on LDLR expression

To explore the mechanisms by which *PROX1* and *CHD7* promote LDL uptake, we next analyzed LDLR protein and mRNA levels in CRISPR-targeted HuH7 cells. *PROX1* targeting led to a significant reduction in surface LDLR receptor expression (26% decrease, 95% CI = 8% to 44%) (Fig 3A), in contrast to *CHD7* targeting which resulted in no significant change (7% decrease, 95% CI = 24% decrease to 10% increase). Quantification of total cellular LDLR protein abundance by immunoblotting (Fig 3B-C) revealed no significant changes in LDLR protein in lysates from *PROX1*-targeted cells (11% reduction, 95% CI = 65% decrease to 43% increase) and *CHD7*-targeted cells (19% increase, 95% CI = 34% decrease to 74% increase). Quantification of *LDLR* mRNA levels by qRT-PCR (Fig 3D) revealed a significant reduction in *LDLR* transcripts upon CRISPR-targeting of *PROX1* (90% reduction, 95% CI = 44% to 100% decrease) but not *CHD7* (33% reduction, 95% CI = 78% decrease to 12% increase). Neither *PROX1* nor *CHD7* targeting were associated with a clear change in LDLR protein half-life (Fig S2A) or protein subcellular localization (Fig S2B). Together with the results of our previously reported CRISPR screen [7], these findings suggest that *PROX1* promotes LDL uptake via positive regulation of *LDLR* gene expression whereas *CHD7* promotes LDL uptake through a LDLR-independent mechanism.

**Figure 3.**
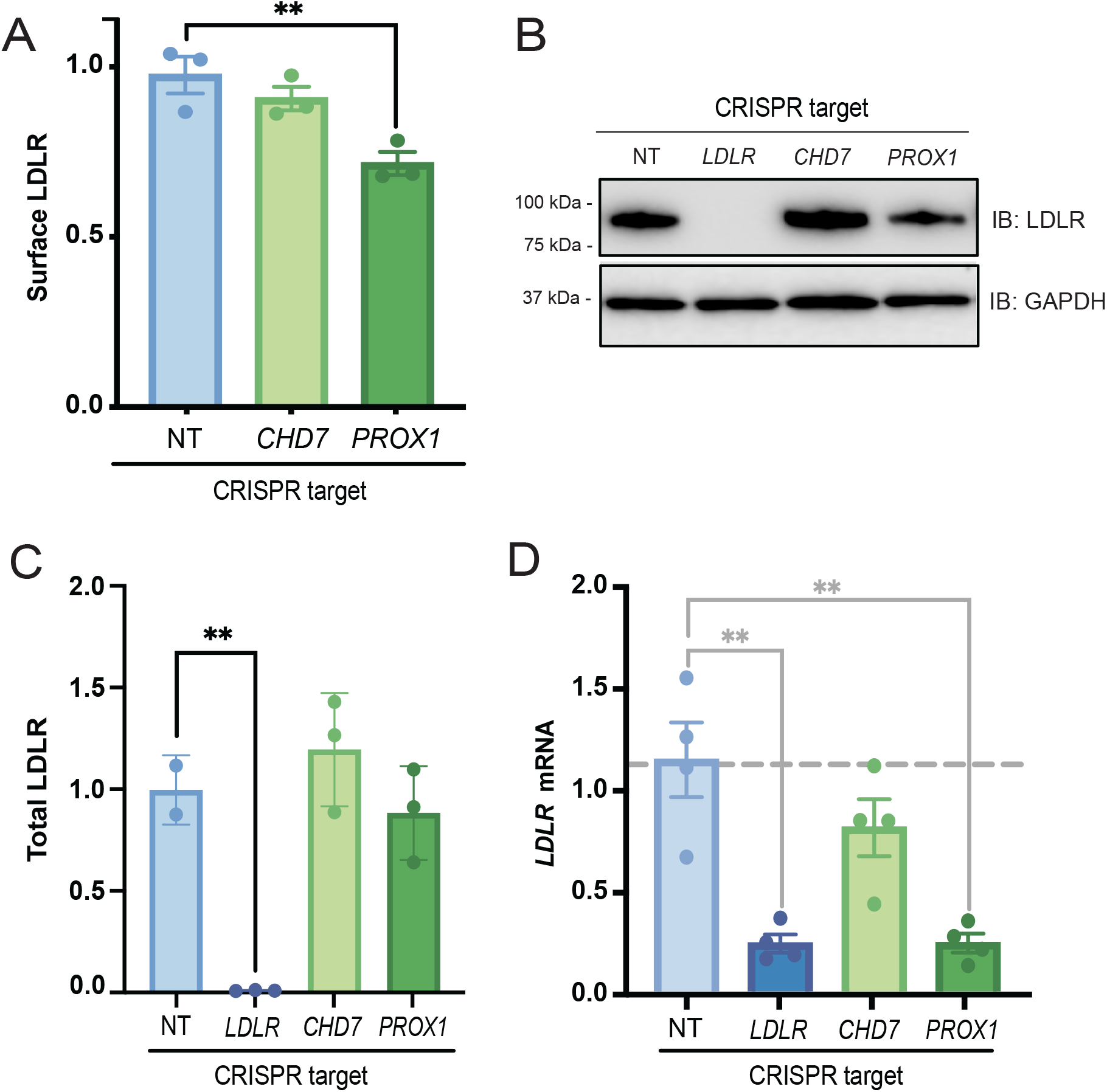
CRISPR targeting of *PROX1*, but not *CHD7*, reduces LDLR expression. HuH7 cells were treated with a control nontargeting (NT) gRNA or gRNAs targeting *LDLR, CHD7*, or *PROX1* and assayed for (A) surface LDLR abundance by flow cytometry analysis, (B-C) total cellular LDLR protein abundance by immunoblotting, and (D) *LDLR* mRNA levels by qRT-PCR. Data points represent independent biological replicates, error bars represent SEM and asterisks indicate p-values by one-way ANOVA (** p < 0.01).

### *Chd7* heterozygous mice exhibit normal plasma cholesterol

Altered cholesterol levels have not been reported previously in patients with CHARGE syndrome. Mice carrying a *Chd7* allele inactivated by a gene trap (Gt) have been previously reported [14], with embryonic lethality observed in homozygous deficient *Chd7^Gt/Gt^* mice by embryonic day 10.5. Heterozygous *Chd7^Gt/+^* mice are viable and recapitulate the developmental abnormalities of CHARGE syndrome. We assayed steady state plasma cholesterol in *Chd7^Gt/+^*and littermate wild type mice and observed no significant differences (Fig 4). These findings are consistent with a lack of effect for a partial reduction in *Chd7* expression on circulating lipoprotein cholesterol levels *in vivo*.

**Figure 4.**
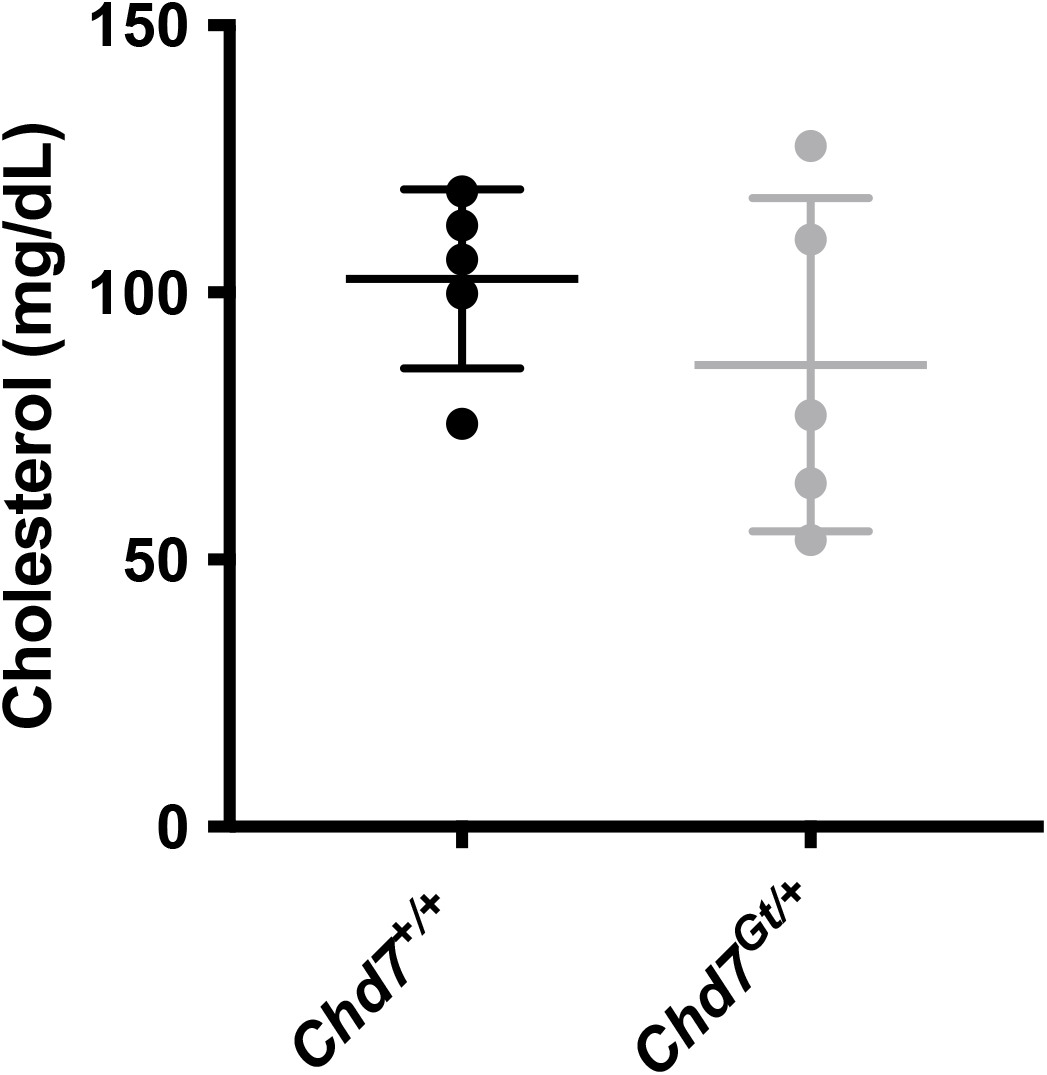
Serum total cholesterol levels are not changed in CHD7-deficient mice. Total cholesterol levels were measured in serum samples obtained from WT ^*Chd7*+/+^ B6129SF1/J mice and heterozygous *Chd7*^*Gt*/+^ deficient mice (n = 5 for each group) (one-way ANOVA).

### Analysis of putative PROX1 and CHD7 binding sites at the *LDLR* locus

We next examined publicly available data to clarify the potential of *PROX1* and *CHD7* to directly regulate hepatic LDLR. Integrating gene expression data from the Genotype-Tissue Expression (GTEx) project [19] and the Broad Dependency Map [20], we observed detectable expression of both genes in RNA-seq data for human liver tissue as well as a number of hepatocyte-derived cell lines, including HuH7, with *CHD7* exhibiting low levels of basal expression in human liver tissue (Fig 5A). *PROX1* mRNA expression was noted to be highest in the liver, among all human tissues tested (Fig 5B).

**Figure 5.**
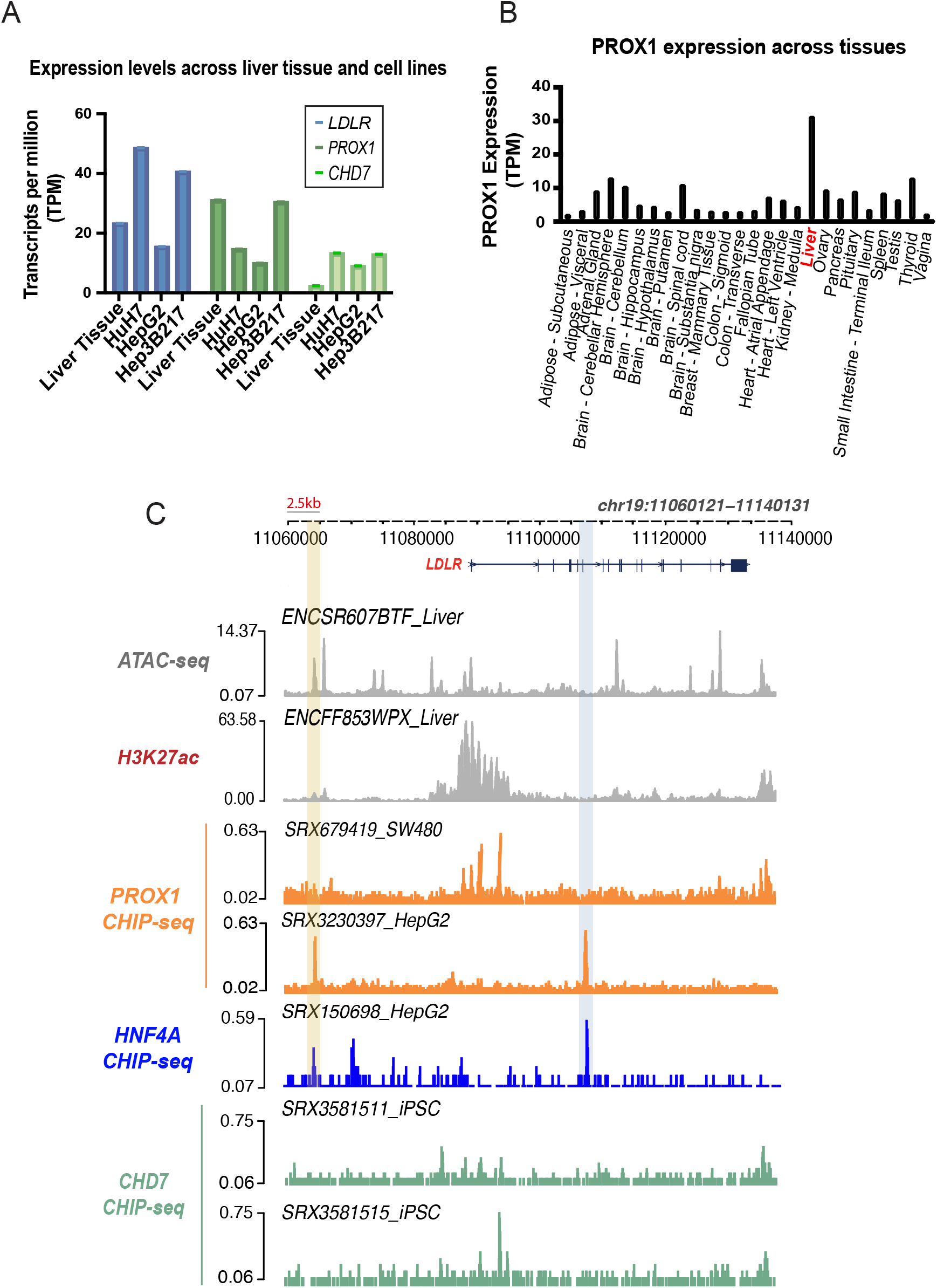
Analysis of *PROX1* and *CHD7* hepatic expression and binding at the *LDLR* locus. **(A)** Transcript levels of *LDLR, PROX1* and *CHD7* in human liver tissue as well as three different liver-derived immortalized cell lines in RNA-seq data in the Broad Dependency Map (DepMap) project. **(B)** *PROX1* transcript levels across a panel of human tissues in the GTEx portal. **(C)** Genomic tracks at the *LDLR* locus of ATAC-seq for the indicated cell line or tissue type, PROX1, HNF4α, and CHD7 ChIP-seq. HNF4α-PROX1 peaks are highlighted in blue and yellow. Data was obtained through the CHIP-atlas and ENCODE project and are identified with experiment unique ID for each. Sources and corresponding references are listed in Methods.

We also analyzed publicly available ChIP-seq data from ChIP-Atlas [21] and ENCODE [22] for evidence of PROX1 or CHD7 binding near *LDLR*. Analysis of PROX1 ChIP-seq data from HepG2 cells revealed two prominent peaks near the *LDLR* locus, one of which colocalized with biochemical features of enhancer activity including open chromatin (as defined by ATAC-seq) and histone H3K27 acetylation (Fig 5C and Fig S3). Both PROX1 ChIP-seq peaks were absent in a similar experiment in a colonic adenocarcinoma cell line (SW480), consistent with cell typespecific binding. Because PROX1 recruitment to genomic binding sites has been reported to involve hepatocyte nuclear factor 4α (HNF4α) [23], and because HNF4α was also identified as a positive regulator of HuH7 LDL uptake in our previously reported CRISPR screen [7], we also analyzed publicly available ChIP-seq data for HNF4α binding at the *LDLR* locus. Both PROX1 ChIP-seq peaks near *LDLR* colocalized with distinct HNF4α ChIP-seq peaks in HepG2 cells. We further analyzed these peaks for PROX1 transcription factor binding motifs using JASPAR [24] and identified at least 3 motifs within these two peaks, with the highest scoring motif falling within the HNF4α-PROX1 binding site that overlapped liver ATAC-seq and H3K27 acetylation ChIP-seq peaks (Fig S4). A similar analysis of CHD7 ChIP-seq data, albeit limited to induced pluripotent stem cells (iPSCs), did not reveal any clear peaks near the *LDLR* locus (Fig 5C). Collectively, these findings are consistent with LDLR-dependent and LDLR-independent mechanisms of LDL uptake regulation by *PROX1* and *CHD7*, respectively.

## Discussion

Given its importance in the pathogenesis of ASCVD, regulation of *LDLR* gene expression has been the subject of extensive investigation. SREBP signaling plays a central role in the regulation of *LDLR* gene expression, with depletion of intracellular sterols triggering the SCAP-mediated trafficking of SREBP and translocation to the nucleus, where SREBP initiates the transcription of lipid metabolism target genes including *LDLR* [8, 25]. The specific binding motifs for SREBP have been defined through comparative analysis of sterol-responsive promoters and mutagenesis experiments of reporter plasmids [26, 27]. Additional transcription factors have been identified that modify *LDLR* expression through sterol-independent mechanisms [28–30]. Unbiased genome-wide CRISPR screens by our group and others have identified other transcriptional regulators whose perturbation influences LDL uptake or LDLR abundance [7, 31].

In this study, we provide functional and bioinformatic evidence of a role for two genes, *PROX1* and *CHD7*, as positive regulators of LDL uptake in HuH7 cells, albeit via distinct mechanisms. *PROX1* acts directly on *LDLR* expression, as supported by the lack of effect of *PROX1* disruption on LDL uptake in *LDLR* knockout cells (Fig S1D), the reduction in *LDLR* mRNA (Fig 3D) and surface protein (Fig 3A) levels caused by *PROX1* disruption, and the identification of PROX1 binding sites (Fig 5C) containing PROX1 binding motifs (Fig S4) near the *LDLR* locus. In contrast, *CHD7* appears to promote LDL uptake independently of LDLR, as indicated by the positive influence of *CHD7* on LDL uptake being unaffected by *LDLR* genetic deletion (Fig S1D), the lack of reduction in *LDLR* mRNA (Fig 3D) or protein (Fig 3A) in *CHD7*-targeted cells, and the lack of CHD7 binding sites near the *LDLR* locus (Fig 5C). Heterozygous *Chd7* knockout mice revealed no change in steady state plasma cholesterol and the embryonic lethality of homozygous knockouts precluded analysis of lipoprotein homeostasis in the complete absence of *Chd7*. Analysis of lipoprotein profiles in CHARGE syndrome patients, heterozygous for *CHD7* loss of function mutations, has not been reported. While *Prox1* genetic ablation in mice also results in embryonic lethality at E14.5 [32], *Prox1* haploinsufficient mice display increased deposition of hepatic lipids [33]. Moreover, liver-specific inactivation of *Prox1* in mice results in hepatic injury and glucose intolerance [34], though an analysis of LDL metabolism in these mice was not reported.

Our findings also demonstrate the confounding influence of HuH7 cell density on quantitative measures of LDLR expression or activity, a finding that is consistent with previous reports [16–18]. We observed a complex effect of cell confluence on LDLR, with increasing density associated with a decrease in LDL uptake and surface LDLR protein abundance but an increase in total cellular LDLR protein and mRNA levels. The underlying mechanism for this effect is unclear but previous hypotheses include cell-cell contact inhibition, restriction of receptor mobility and/or inhibition of bulk endocytosis caused by confluent monolayer formation [16–18]. Regardless of the mechanism for these effects, we successfully reduced experimental variation by carefully accounting for cell density and normalizing readouts to a standard curve with a range of densities for control cells. Application of the approach reported here may improve the power to detect changes in LDLR expression in cultured cells for future studies.

Other groups have reported an association of *PROX1* levels with LDLR expression, but in conflicting directions to what we observed here. Overexpression of *PROX1* has been reported to decrease *LDLR* mRNA levels in endothelial cells [35], while an RNA interference screen of HuH7 cells found *PROX1* targeting to confer an increase in LDL uptake [36]. The basis for these discrepancies is not clear, but may be related to differences in cell type, culture conditions, or mechanism of genetic perturbation. The latter study, for example, did not detect an influence for *LDLR* itself, or for the canonical regulators *SCAP, MBTPS1*, or *MBTPS2* [38], suggesting that the conditions of the screen may not have been optimal for uncovering functional modifiers of LDLR expression. Others have provided evidence that *PROX1* has critical functions in adult liver lipid homeostasis, with *PROX1* disruption leading to metabolic transcriptional changes, upregulation of hepatic lipid deposition, and obesity [33, 34, 37]. *PROX1* has also been reported to be recruited for transcriptional regulation of lipid metabolism in mouse liver tissue by HNF4α, [23, 38], another transcription factor with an established role in liver development and lipid homeostasis [39–42]. The overlap in PROX1 and HFN4α ChIP-seq peaks in HepG2 cells suggests that these transcription factors may be co-recruited to promote expression of hepatic *LDLR*.

Overall, our findings build upon our prior CRISPR screen to validate two functional regulators of HuH7 cellular LDL uptake and clarify their mechanisms of action [7]. Our findings are consistent with *PROX1* acting as a liver-specific direct positive regulator of *LDLR* gene expression and *CHD7* promoting LDL uptake though a LDLR-independent mechanism. Future studies will be necessary to establish the physiologic relevance of these findings *in vivo* and their potential as therapeutic targets for the prevention and treatment of ASCVD.

## Materials and Methods

### Reagents

HuH7 cells were cultured in Dulbecco’s Modified Eagle Medium (DMEM) (ThermoFisher Scientific) supplemented with 10% fetal bovine serum (FBS, Sigma) and 100 U/ml penicillin, and 100 mg/ml streptomycin (ThermoFisher Scientific) at 37°C in a 5% CO2-conditioned humidified incubator. CRISPR-mediated gene disruption was performed by identifying individual gRNA sequences with consistent enrichment in LDL^low^ cells in our previously reported CRISPR screens [7] and cloning these sequences into the BsmBI sites of pLentiCRISPRv2 (Addgene #52961, [43]) or into the BbsI sites of pX459 (Addgene #62988, [44]). Lentiviral particles were then prepared by cotransfection of pLentiCRISPRv2 constructs into HEK293T cells with pCMV-VSV-G (Addgene #8454, a gift from Bob Weinberg) and psPAX2 (Addgene #12260, a gift from Didier Trono) using Lipofectamine LTX (ThermoFisher Scientific) as previously reported [7]. Conditioned media containing virus was harvested at 48 hours post-transfection and single use aliquots were snap frozen in liquid nitrogen and stored at −80°C. To generate pooled CRISPR-targeted lines, HuH7 cells were transduced with lentivirus, selected with 3μg/mL puromycin, and passaged for two weeks in order to allow target site mutagenesis and turnover of wild-type protein.

### Flow cytometry analysis for LDL uptake and surface LDLR

Control and CRISPR-edited cells were seeded into 6-well plates at a range of densities. Two days later, CRISPR-edited cells were selected from wells at ~60-90% confluence while control cells were analyzed at a range spanning from ~30-40% confluence to fully confluent monolayers. To quantify LDL uptake, cells in selected wells were first washed with pre-warmed serum-free DMEM and then incubated in serum free DMEM containing 4 μg/mL fluorescently DyLight^550^-conjugated LDL (Cayman Chemical) at 37°C for 1 hour. Cells were then washed with blocking buffer (PBS + 2% FBS), harvested with TrypLE Express, washed three times with blocking buffer, stained with Sytox Blue (ThermoFisher Scientific) viability dye, and resuspended in PBS containing a known concentration of CountBright Absolute Counting Beads (ThermoFisher Scientific). Total cell numbers in each well were then calculated by adjusting the observed concentration of cell events to bead events during flow cytometry. The mean fluorescent intensity (MFI) of internalized LDL was plotted against the calculated total cell number across plating densities and a linear regression equation was used to calculate the expected LDL fluorescence for a given cell number. Experimental samples were then calculated based on the observed LDL MFI relative to this predicted MFI. For surface LDL receptor staining, HuH7 cells were detached with TrypLE Express, washed three times with ice cold blocking buffer, resuspended at approximately 10^6 cells/mL in blocking buffer and incubated for 30 minutes with end-over-end rotation at 4°C. After centrifugation at 500g, cells were resuspended with fluorescently labelled LDLR antibody in blocking buffer and incubated for 1 hour in the dark at 4°C. Cells were washed with cold PBS three times, resuspended in 200 μl PBS, and incubated with Sytox Blue prior to analysis by flow cytometry. Data analysis was performed using FlowJo and GraphPad Prism.

### Western blot

HuH7 cells were collected with TrypLE express, washed in PBS, then lysed in RIPA lysis and extraction buffer (ThermoFisher Scientific) containing complete protease inhibitor cocktail (Roche). Lysed cell suspensions were rotated in 4°C for 1 hour, followed by centrifugation at 15000g for 1 hour at 4°C and collection of supernatants. Protein concentration was determined with the Bio-Rad DC assay (Bio-Rad) and 10 μg of protein lysate was fractionated on a NuPAGE 4-12%, Bis-Tris mini protein gel (Thermo Scientific) according to manufacturer’s instructions. Separated proteins were then transferred to nitrocellulose membranes (Thermo Scientific) using the iBlot 2 Dry Blotting System (Thermo Scientific) at 20V for 7 minutes. *CHD7* blots were transferred at 12V for 18 hours. Membranes were blocked with 5% non-fat milk in Tris-buffered saline with 0.1% Tween (TBST) overnight, probed with primary and secondary antibodies in blocking buffer for 1 hour, washed with TBST and developed using SuperSignal West Femto Maximum Sensitivity Substrate (Thermo Scientific). Analysis of Western blots was performed using Bio-Rad Image Lab Software (version 6.1, Image Lab) with normalization of protein abundance to GAPDH.

### Immunofluorescence and Confocal Microscopy

Cells were plated on poly-D-lysine coated glass coverslips (Electron Microscopy Sciences) and fixed with 4% paraformaldehyde for 20 minutes at room temperature. After washing three times with PBS, cells were incubated at room temperature for 1 hour in blocking buffer (5% FBS, 50 mM glycine in PBS, supplemented with 0.75% TritonX-100), then incubated with primary antibodies for 1 hour, washed thrice, stained with secondary antibody for another hour and washed five times prior to mounting. Coverslips were mounted on glass slides with Prolong Diamond mounting media (Fisher, #P36965). Images were acquired with a NIKON A1 standard sensitivity (SS) confocal microscope with 60X objective (NA51.4) and image analysis was performed using ImageJ/Fiji [45, 46].

### Analysis of Gene Expression

For qRT-PCR, total RNA was extracted with a RNeasy kit (Qiagen) and first strand cDNA prepared using the PrimeScript RT (Clontech) according to manufacturer’s instructions. Amplification was performed using Power SYBR Green PCR Master Mix (Thermo Fisher) and analyzed using the QuantStudio 5 Real-Time PCR System. Relative *LDLR* transcript levels were normalized against the mean C_t_ value for the control genes *RPL37* and *RPL38*. Total cell numbers in each analyzed well were quantified in parallel by flow cytometry with comparison to a standard curve of control cells at a range of densities, as described above.

### Quantification of LDLR protein turnover

To assess LDLR protein stability, control and CRISPR-targeted HuH7 cells were incubated at 37°C with 10 μg/mL cycloheximide, and samples were collected at serial time points (0, 2, 4, 6 and 8 hours post-treatment). Cells were fixed and total cellular LDLR protein abundance was quantified by flow cytometry, as described above.

### Mouse phenotyping

Following euthanasia by isoflurane inhalation, blood was drawn from the inferior vena cava of *Chd7^+/+^* and *Chd7^Gt/+^* mice (see Key resource table) using a 23 G needle and a 1 ml syringe.Blood was collected into a serum separator tube (365967, BD, Franklin Lakes NJ), allowed to clot at room temperature for at least 10 minutes, and centrifuged at 15,000 g for 10 minutes to separate serum. Sera were aliquoted and stored at −80°C. Sera were analyzed by a colorimetric assay for total cholesterol (SB-1010-225, Fisher Scientific, Hampton NH) as previously described [7]. All animal care and use complied with the Principles of Laboratory and Animal Care established by the National Society for Medical Research. All animal protocols in this study were approved by the Institutional Animal Care and Use Committee (IACUC) of the University of Michigan (protocol number PRO00009304). Both male and female mice were used in this study.

### Bioinformatic analysis

*PROX1* and *CHD7* expression levels for human cell lines were obtained from the Broad Dependency Map Portal [20] (DepMap Public 22Q2). PROX1 expression levels in human tissues were obtained from the Genotype-Tissue Expression (GTEx) database [19]. CHIP-seq and ATAC-seq data sets were downloaded from the ChIP-Atlas [21] or ENCODE [47] databases: All ENCODE experiments of ATAC-seq or H3K27ac ChIP-seq of liver tissue that had warnings for low read counts or coverage were analyzed (ID: ENCSR607BTF, ENCFF831BRS, ENCFF646EEM, ENCFF556YQA, ENCFF241ZLC, ENCFF469MOB and ENCFF232QBB), HNF4α CHIP-seq (ID: SRX150698), PROX1 HepG2 CHIP-seq (ID: SRX3230397), PROX1 SW480 CHIP-seq (ID: SRX679419), CHD7 iPSC CHIP-seq (ID: SRX3581511 and SRX3581515), Liver tissue H3K27ac CHIP-seq (ID: ENCFF352AAW, ENCFF275OLK and ENCFF853WPX). Identifiers are specific to the source (i.e., EN for ENCODE datasets and ERX/SRX for CHIP-atlas). Peaks were visualized with the *trackplot* (github.com/PoisonAlien/trackplot/) R package. We calculated relative scores for putative PROX1 binding sites scores across chromosome 19 using the JASPAR Position frequency matrices (PFMs) corresponding to JASPAR PROX1 binding motif (MA0794.1) with an 80% relative score cutoff using the ‘searchSeq’ function in TBFS Tools R package. Motifs of interest were identified within the chromosome coordinates corresponding to the HNF4α-PROX1 overlapping CHIP-seq peaks.

#### Key Resources Table

| Key Resources Table |  |  |  |
| --- | --- | --- | --- |
| Reagent Type or Resource | ID | Source or Reference | Comments |
| Antibody | LDL Receptor | Abcam; ab52818 | Western Blot (1:1000) |
| Antibody | LDL Receptor | R&D system; FAB2148G Alexa488 | FACS (5ul/100ul (10<sup>6</sup> cells)) |
| Antibody | GAPDH | Abcam; ab181602 | Western Blot (1:5000) |
| Antibody | CHD7 | Cell Signaling Technologies; 6505S | Western Blot (1:1000) |
| Antibody | PROX1 | R&D Systems; AF2727SP | Western Blot (1:200) |
| Antibody | Goat Anti-Rabbit IgG (H + L)-HRP Conjugate | Bio-Rad (#1706515) | Western Blot (1:5000) |
| Reagent | LDL-Dylight550 | Cayman Chemical (10011229) | FACS (4ug/mL) |
| Reagent | Sytox-Blue | Fisher (S34857) | FACS |
| Reagent | CountBright™ Absolute Counting Beads | Thermo Scientific (C36950) | FACS |
| Organism | Mouse Strain B6; 129S- *Chd7*<sup>Gt(S20-7E1)</sup><sub>Sor</sub>/*DmmJ* | The Jackson Laboratory | Strain #030659 [14] |
| Software | Prism 7 for Mac OS X GraphPad | https://www.graphpad.com/scientific-software/prism/ |  |
| Software | R | R Development Core Team, 2016 | https://www.r-project.org |
| Software | Integrative Genome Browser | https://software.broadinstitute.org/software/igv/ | [48] |
| Software | Adobe Illustrator | Adobe Inc. (2022) | https://adobe.com/products/illustrator |
| Software | FIJI | [49] |  |
| Database | ENCODE Consortium | https://www.encodeproject.org/ |  |
| Database | CHIP-Atlas | https://chip-atlas.org/ |  |
| Database | Broad Dependency Map Portal (DepMap) | https://depmap.org/portal/ | [20] |
| Database | The Genotype-Tissue Expression (GTEx) project | https://gtexportal.org/home/ | [19] |
| Algorithm | JASPAR 2020 | http://bioconductor.org/packages/release/data/annotation/html/JASPAR2020.html |  |
| Algorithm | TFBSTools | https://bioconductor.org/packages/release/bioc/html/TFBSTools.html |  |
| Algorithm | Biostrings | https://bioconductor.org/packages/release/bioc/html/Biostrings.html |  |
| Algorithm | *trackplot* | https://www.biostars.org/p/475853/ |  |
| Algorithm | *karyoploteR* | https://bernatgel.github.io/karyoploter_tutorial/ |  |

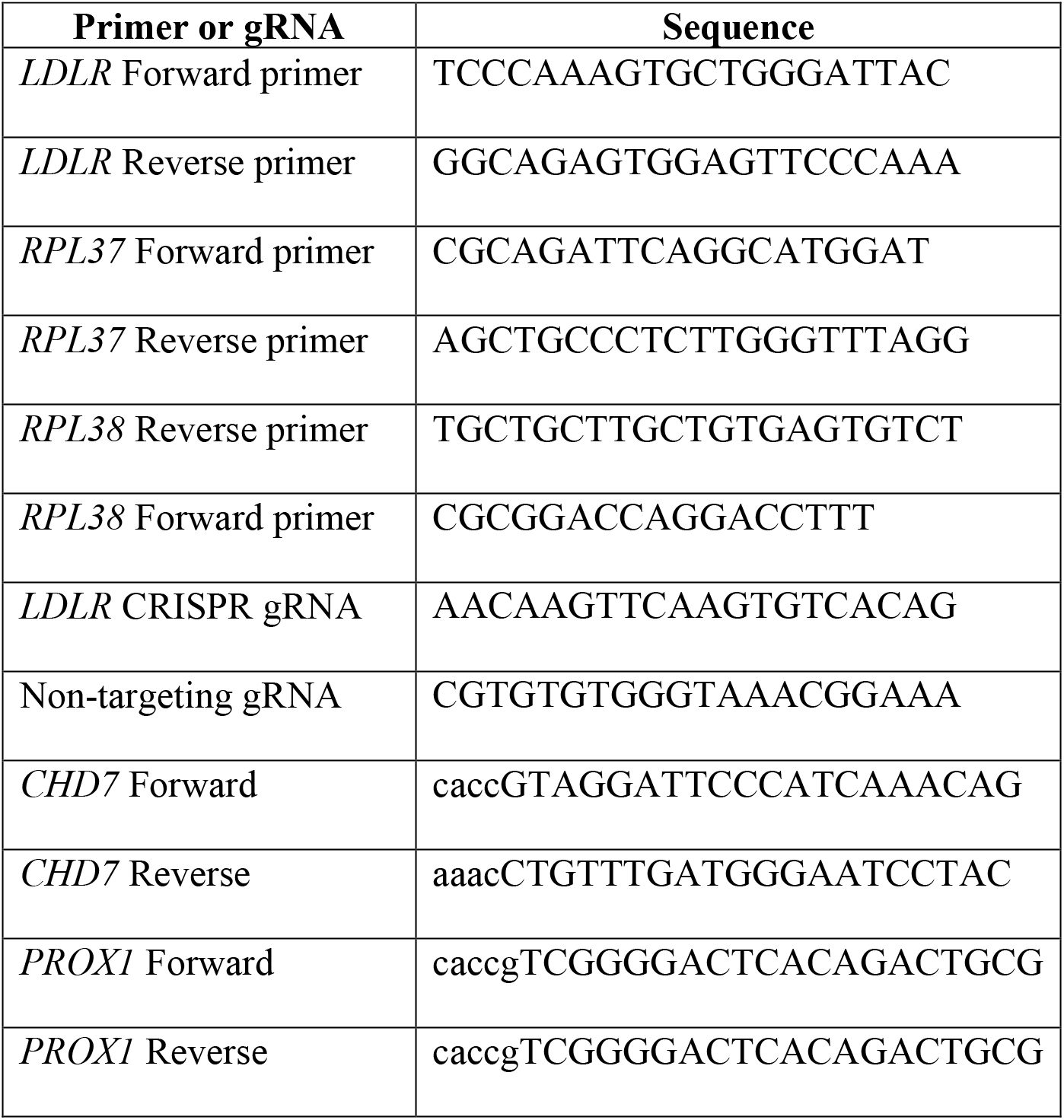

## Acknowledgments

This work is supported by the HHMI Gilliam Fellowship for Advanced Study (C. S. Z.), NIH grants R35-HL135793T (D. G.), K08-HL148552 (B. T. E.), R01-DC018404 (D. M. M.), the Taubman Medical Institute (D. M. M.), and the Ravitz Foundation Professorship in Pediatrics and Communicable Diseases (D. M. M.). D.G. is a Howard Hughes Medical Institute Investigator.

## Notes

### Competing Interest Statement

The authors have declared no competing interest.

