## Supplementary Figures for "Regulation of cellular LDL uptake by *PROX1* and *CHD7*"

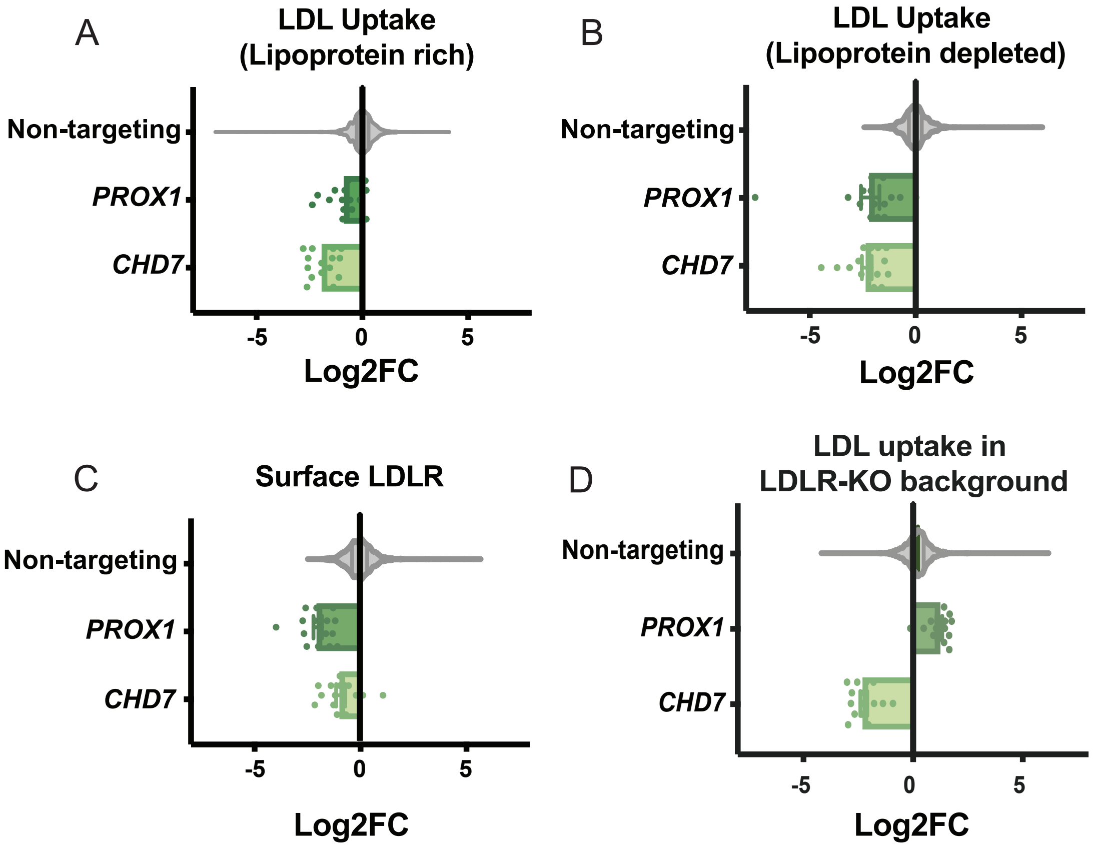


**Supplementary Figure 1 – Enrichment analysis for individual gRNAs targeting *PROX1* and *CHD7* in data from previous CRISPR screen.** Log2 fold-change (Log2FC) for each of 15 individual gRNAs targeting *PROX1* or *CHD7* are plotted relative to the distribution of values for 1000 control non-targeting gRNAs for each independent CRISPR screen, including (A) HuH7 cells grown in lipoprotein rich media sorted based on their relative uptake of LDL (negative value indicates gene disruption reduces LDL uptake); (B) HuH7 cells preincubated in lipoprotein-deficient media to stimulate LDLR expression and then sorted based on their relative uptake of LDL (negative value indicates gene disruption reduces LDL uptake); (C) HuH7 cells sorted based on their surface LDLR abundance (negative value indicates gene disruption reduces surface LDLR abundance); and (D) *LDLR* knockout cells sorted based on their relative uptake of LDL (negative value indicates gene disruption reduces LDL uptake).

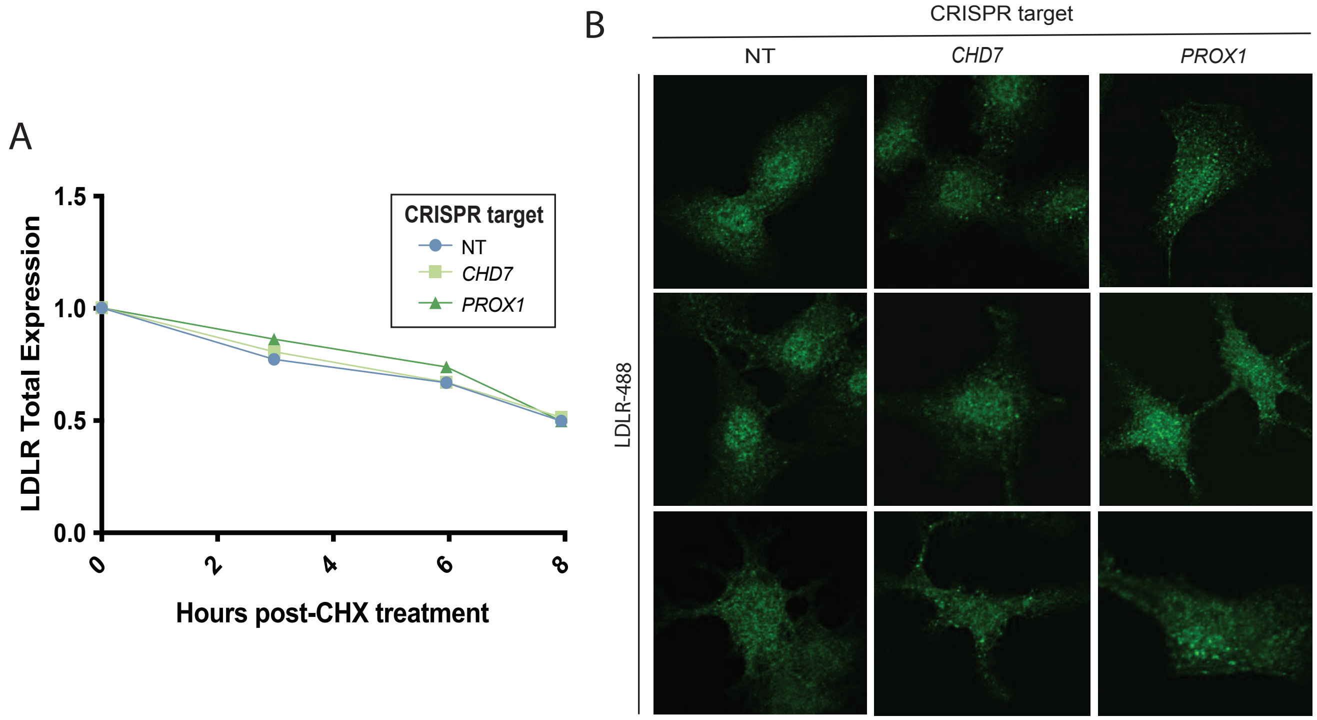


**Supplementary Figure 2 – CRISPR-targeting of *CHD7* or *PROX1* does not change LDLR protein stability (A)** LDLR degradation rate in HuH7 cells treated with CRISPR and *CHD7* or *PROX1* gRNAs were compared to an NT gRNA at several timepoints after addition of cycloheximide (10µm; CHX) **(B)** Immunofluorescence for LDLR in HuH7 cells treated with a NT gRNA, or gRNAs targeting *CHD7* or *PROX1*.

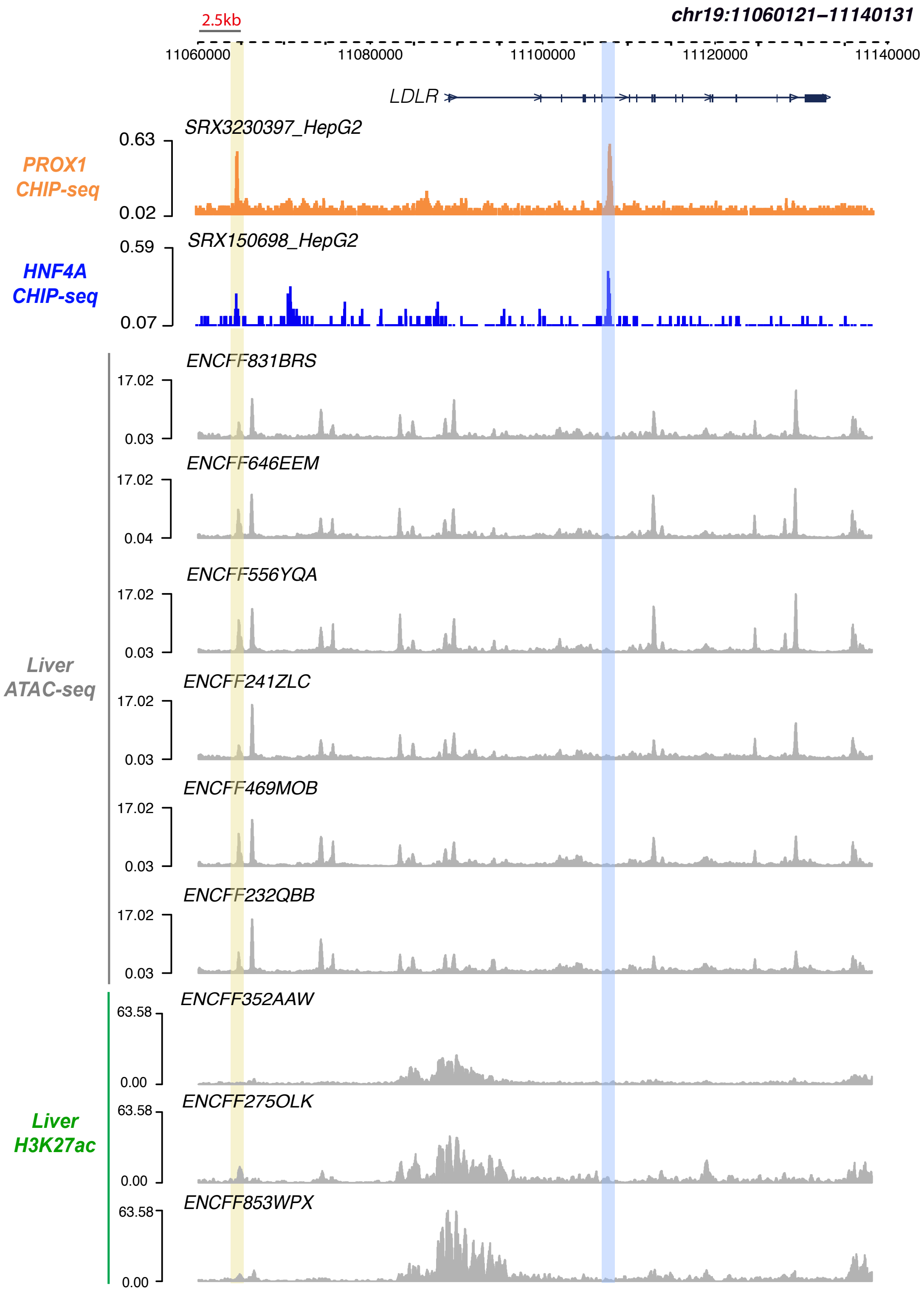


**Supplementary Figure 3 – Colocalization of PROX1 binding sites at the *LDLR* locus with HNF4α binding sites and biochemical features of enhancer activity in liver tissue.** The genomic track for HepG2 PROX1 ChIP-seq is displayed in parallel with tracks for HepG2 HNF4α ChIP-seq and for liver tissue ATAC-seq or H3K27ac-seq experiments. Colocalized binding peaks with HNF4α binding are highlighted with (yellow) or without (blue) association with liver tissue ATAC-seq and H3K27ac ChIP-seq peaks.

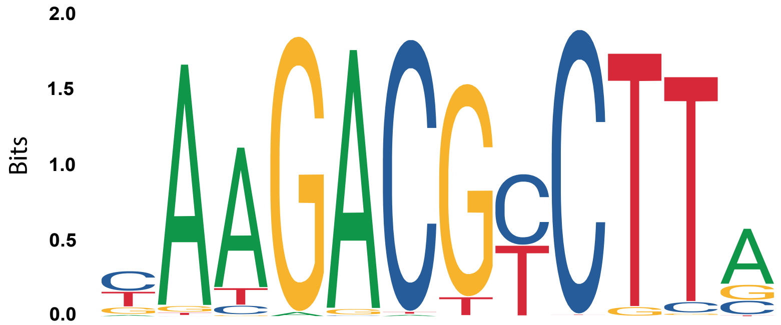


| Genomic coordinates | 1 | 2 | 3 | 4 | 5 | 6 | 7 | 8 | 9 | 10 | 11 | 12 | Rel. Score | p-value |
|---|---|---|---|---|---|---|---|---|---|---|---|---|---|---|
| chr19:11064833- 11064844 (+ strand) | G | G | T | G | A | C | T | C | C | T | T | C | 0.86 | 0.0002 |
| chr19:11108317-11108328 (- strand) | C | A | A | G | A | C | T | C | T | G | T | T | 0.81 | 0.0013 |
| chr19:11108258 -11108269 (+ strand) | C | A | T | G | A | A | T | T | C | T | T | T | 0.83 | 0.0006 |

**Supplementary Figure 4 – PROX1 Binding motif sequence similarity at HNF4α-PROX1 overlapping peaks near the *LDLR* locus.** The consensus PROX1 binding motif (JASPAR 2022 MA0794.1) is displayed in alignment with statistically significant (p-value< 0.01) sequences identified in the 2 HNF4α-PROX1 binding peaks near the *LDLR* locus. Sequences matching the top-scoring nucleotide in the consensus motif are highlight in yellow.
